# A novel SARS-CoV-2 related coronavirus in bats from Cambodia

**DOI:** 10.1101/2021.01.26.428212

**Authors:** Vibol Hul, Deborah Delaune, Erik A Karlsson, Alexandre Hassanin, Putita Ou Tey, Artem Baidaliuk, Fabiana Gámbaro, Vuong Tan Tu, Lucy Keatts, Jonna Mazet, Christine Johnson, Philippe Buchy, Philippe Dussart, Tracey Goldstein, Etienne Simon-Lorière, Veasna Duong

## Abstract

Knowledge of the origin and reservoir of the coronavirus responsible for the ongoing COVID-19 pandemic is still fragmentary. To date, the closest relatives to SARS-CoV-2 have been detected in *Rhinolophus* bats sampled in the Yunnan province, China. Here we describe the identification of SARS-CoV-2 related coronaviruses in two *Rhinolophus shameli* bats sampled in Cambodia in 2010. Metagenomic sequencing identified nearly identical viruses sharing 92.6% nucleotide identity with SARS-CoV-2. Most genomic regions are closely related to SARS-CoV-2, with the exception of a small region corresponding to the spike N terminal domain. The discovery of these viruses in a bat species not found in China indicates that SARS-CoV-2 related viruses have a much wider geographic distribution than previously understood, and suggests that Southeast Asia represents a key area to consider in the ongoing search for the origins of SARS-CoV-2, and in future surveillance for coronaviruses.

## Main Text

Over a year has passed since the emergence of Severe Acute Respiratory Syndrome coronavirus 2 (SARS-CoV-2)^1^, responsible for the ongoing coronavirus disease 2019 (COVID-19) pandemic. However, information on the origin, reservoir, diversity, and extent of circulation of ancestors to SARS-CoV-2 remains scarce. Horseshoe bats (genus *Rhinolophus*) are believed to be the main natural reservoir of SARS-related coronaviruses also named Sarbecoviruses^2^. Indeed, a high diversity of coronavirus species have been found in *Rhinolophus* bats collected in several provinces of China^3^. To date, the closest relatives to SARS-CoV-2 were identified from bats sampled in the Yunnan province, southern China^1,4^. RaTG13 was isolated from a *Rhinolophus affinis* horseshoe bat in 2013, and RmYN02 from a *Rhinolophus malayanus bats* in 2019. Two viruses were also detected in Sunda pangolins (*Manis javanica*) seized in two provinces of southern China^5^. More distant and highly mosaic recombinant viruses were also sampled from bats in the Zhejiang province, in eastern China in 2015 and 2017.

Southeast Asia is considered a hotspot for emerging diseases^6^, and more than 25% of the worlds bat diversity is found in there^7^. Following the emergence of COVID-19, to search for putative SARS-CoV-2-like betacoronaviruses (betaCoVs), 430 archived samples from 6 bat families and 2 carnivorous mammal families, including 162 oral swabs and 268 rectal swabs, were retrospectively tested with a pan-coronavirus (pan-CoV) hemi-nested RT-PCR^8^ (Supplementary table 1). Sixteen out of 430 (3.72%) samples tested positive for CoV by pan-CoV hemi-nested PCR. Eleven were classified as alphacoronaviruses and 5 as betaCoV. Two of the 5 betaCoV samples further tested positive using a RT-qPCR targeting the RdRp gene of sarbecoviruses^9^. Both samples came from Shamel’s horseshoe bats (*Rhinolophus shameli*) sampled in December 2010 in the Steung Treng province in northeastern Cambodia.

RNA samples were then processed for next-generation metagenomic sequencing, using a ribosomal RNA depletion approach and randomly primed cDNA synthesis^10^. Read assembly reconstructed two nearly identical near full-length coronavirus genomes, named BetaCoV/ Cambodia/RShSTT182/2010 (RshSTT182) and BetaCoV/Cambodia/RShSTT200/2010 (RshSTT200), respectively. The two sequences are closely related to SARS-CoV-2, exhibiting 92.6% nucleotide identity across the genome (Supplementary table 2) and identical genomic organization. Phylogenetic analysis using full genome sequences shows that RshSTT182 and RshSTT200 represent a new sublineage of SARS-CoV-2 related viruses, despite the geographic distance of isolation (Figure 1). Genetic similarity with SARS-CoV-2 is maintained across the genome, with the exception of a portion corresponding to the spike N terminal domain (NTD; Figure 2 and suppl 2). In several sections of the genome, including the region spanning nsp4 to nsp8 within orf1a, and orf8, RshSTT182 and RshSTT200 are genetically closer to SARS-CoV-2 than the any other closely related viruses discovered to date. Similarity is further evidenced when inferring phylogeny based on the sequence coding for these proteins.

**Figure 1.**
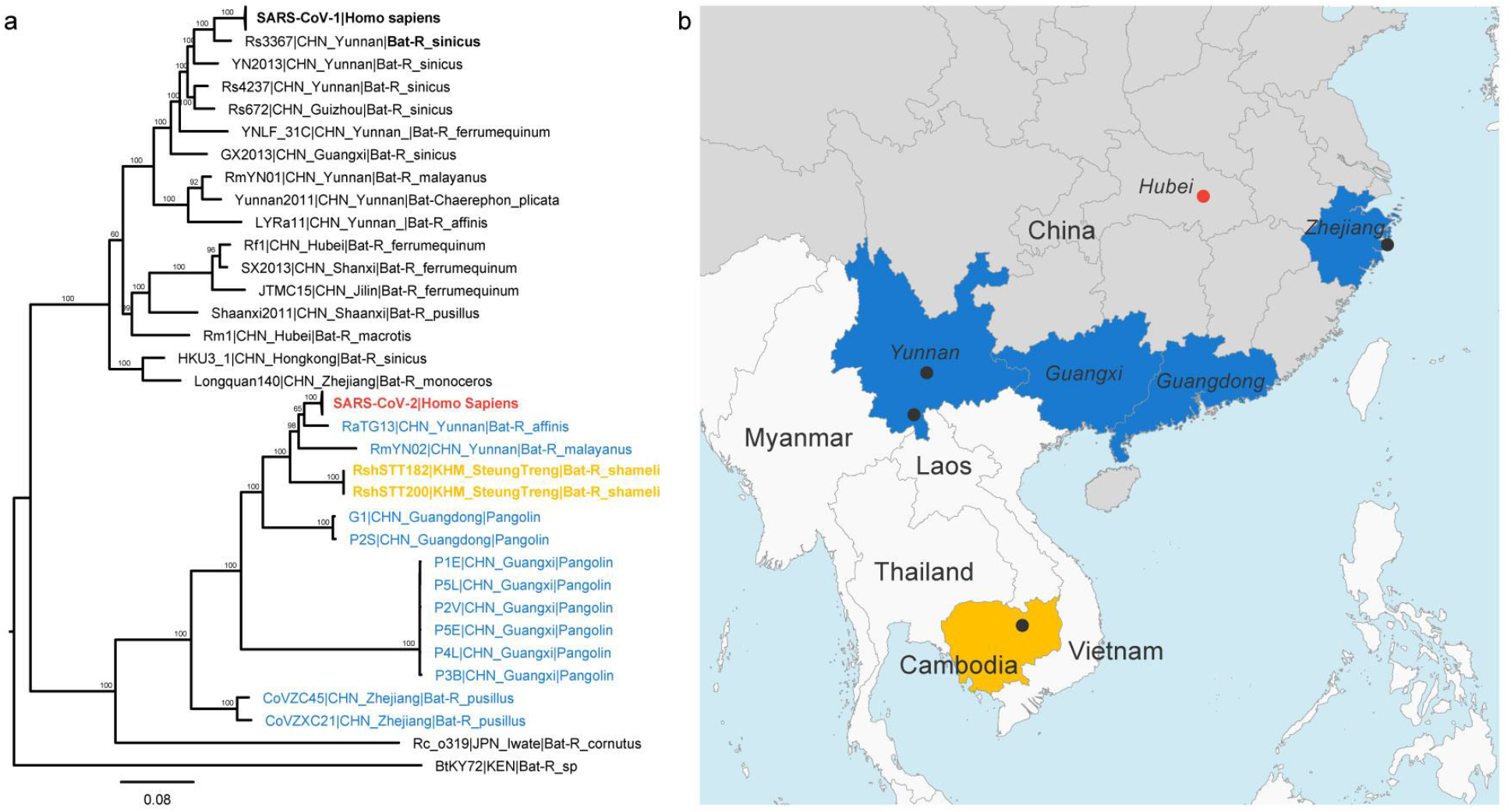
Phylogenetic analysis of SARS-CoV-2 and representative sarbecoviruses and geographical context. a. Maximum likelihood phylogeny of the subgenus Sarbecovirus (genus *Betacoronavirus*; n = 37) estimated from complete genome sequences using IQ-TREE and 1,000 replicates. The coronaviruses of the SARS-CoV-2 lineage are color coded by country of sampling as on the map. In orange, Cambodia, and shades of blue, China. Taxa names include the isolate name, country and province of sampling, and host. The scientific names of the hosts are abbreviated as follows: Bats: *R. affinis, Rhinolophus affinis; R. sinicus, Rhinolophus sinicus; R. ferrumequinum, Rhinolophus ferrumequinum; R. malayanus, Rhinolophus malayanus; C. plicata, Chaerephon plicata; R. pusillus, Rhinolophus pusillus; R. macrotis, Rhinolophus macrotis; R. monoceros, Rhinolophus monoceros; R. cornutus, Rhinolophus cornutus; Pangolin: M_javanica, Manis javanica* and human: H. sapiens, *Homo sapiens*. A maximum clade credibility tree is available in Supplementary Figure 3. b. map of parts of China and Southeast Asia. Regions where viruses of the SARS-CoV-2 lineage were sampled are colored as in the tree. A black dot indicates a sampling site when known, and the red dot shows the location of Wuhan, where the first cases of SARS-CoV-2 infection were reported.

**Figure 2.**
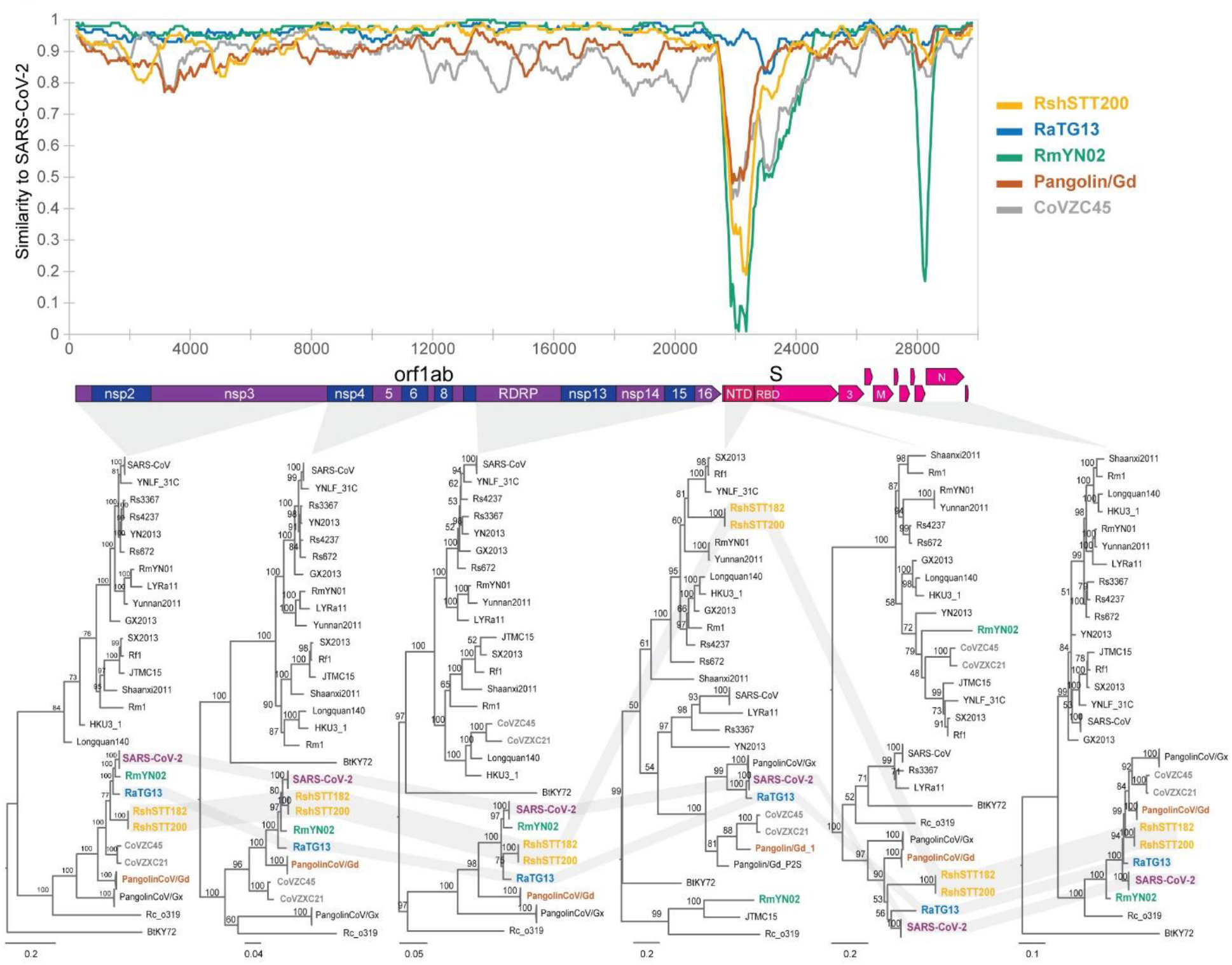
Recombination analysis. a. Sliding window analysis of changing patterns of sequence similarity between SARS-CoV-2 and related coronaviruses from China and Cambodia. ZXC21 and ZC45 were merged for this analysis. b. Phylogenetic tree of different genomic regions. Branch support obtained from 1,000 bootstrap replicates are shown. SARS-CoV and SARS-CoV-2 sequences are collapsed, and trees are midpoint rooted for clarity.

Extensive evidence exists on numerous recombination events in the evolutionary history of the sarbecoviruses^11-13^. Consistent with this, we found that both RshSTT182 and RshSTT200 are also mosaic viruses (Figure 2 and supplementary figure 2); however, most regions identified as recombinant in origin appear to have involved close relatives within the SARS-CoV-2 sublineage.

Only a region encompassing the Spike N terminal domain (NTD) is closer to more distantly related betaCoVs. In all other regions of the genome, the viruses detected in Cambodia consistently branch as a sister clade to SARS-CoV-2 and RaTG13, with minor swaps in the subtree topology. Interestingly, both regions showing high similarity to SARS-CoV02 (nsp4 to 8 within orf1a and orf8) overlap with regions identified as recombinant. All these elements suggest a co-circulation of ancestors to these viral sublineages with both a wider geographic area and more distinct bat species than those previously identified. Of note, the current geographic distribution of *R. shameli* bats does not include China (Supplementary Figure 4)^14^. However, the distribution of *R. affinis* and *R. malayanus* overlap with *R. shameli* distribution area in Southeast Asia, and extend into China, including the Yunnan province where the other two viruses closely related to SARS-CoV-2 were sampled. In addition, the haplotype network of *R. shameli* CO1 sequences shows a typical star-like pattern, suggesting that populations of *R. shameli* found between northern Cambodia and northern Laos are not genetically isolated. Some bat species are believed to be migratory, and distinct species of bats often share habitats, and even co-roost, if only transiently. Finally, *R. affinis* and *R. malayanus* bats were concomitantly captured in the same northeastern karst region where these *R. shameli* bats were sampled.

Further risk assessment is needed to understand the host range (including Humans) and pathogenesis associated with this novel sublineage. Homology modeling suggests that the external subdomain of the spike receptor binding domain (RBD) structure is highly similar to SARS-CoV-2 (Figure 3). We note the shortening of a loop at the beginning of the receptor binding motif and the presence of a conserved disulfide bond. Importantly, five of the six amino acid residues reported to be major determinants of efficient receptor binding of SARS-CoV-2 to the human angiotensin-converting enzyme 2 (ACE2) receptor ^15^ are conserved. Finally, the poly-basic (furin) site present in SARS-CoV-2 is absent in both RshSTT182 and RshSTT200.

**Figure 3.**
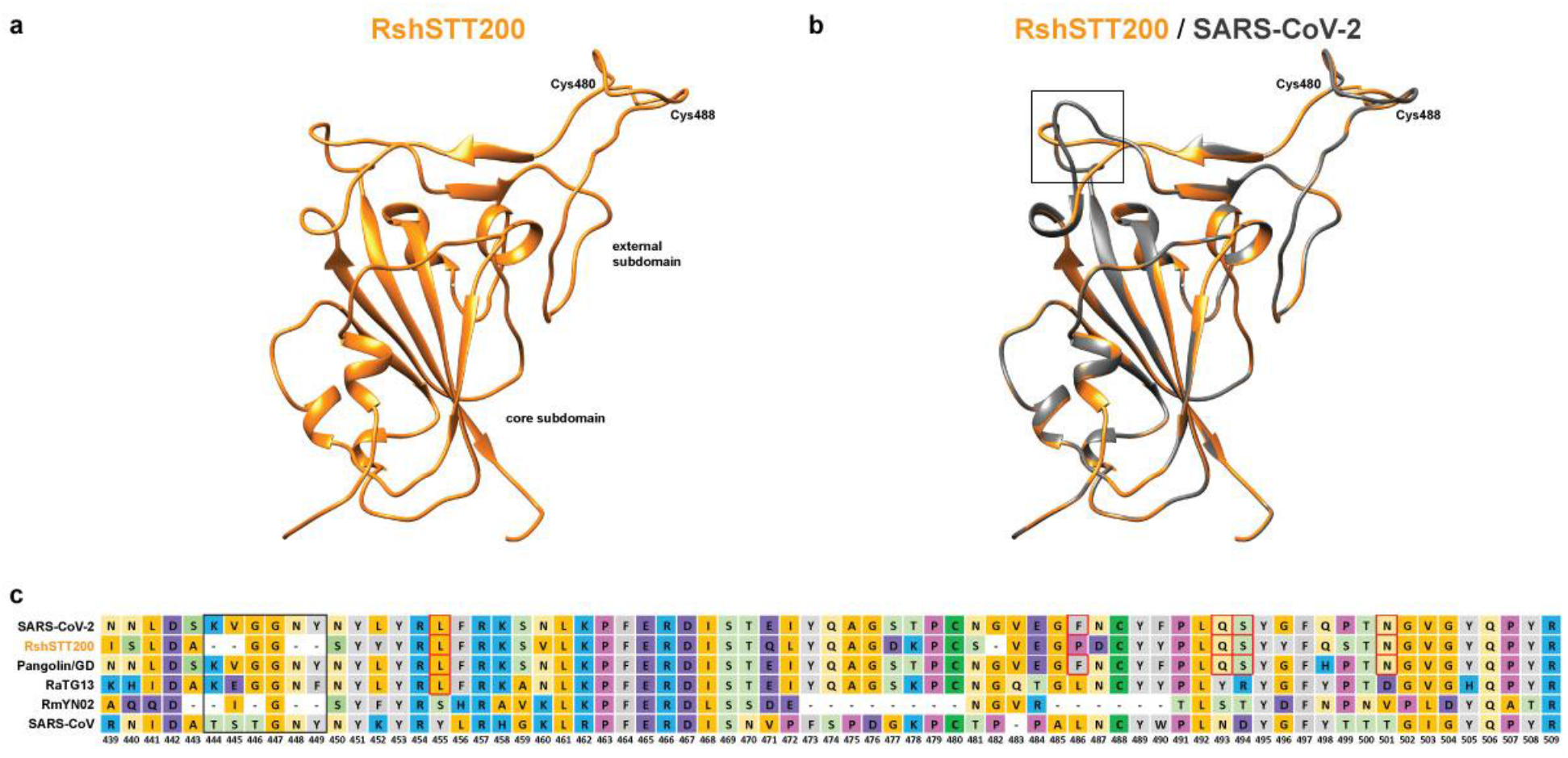
Structural modeling a. Homology modeling of the RBD structure. The three-dimensional structure of the RshSTT200 Spike RBD was modeled using the Swiss-Model program employing the structure of SARS-CoV-2 (PDB: 6yla.1) as a template. The core and external subdomains are colored orange, and grey for RshSTT200 and SARS-CoV-2, respectively. The shortening a loop near the receptor-binding site of RshSTT200 are indicated by a black rectangle. The cysteines involved in a conserved disulfide bond are indicated. b. Alignment of the RBM amino acid sequences of selected betaCoVs.

The data presented here indicates that SARS-CoV-2 related viruses have a much wider geographic distribution than previously understood, and likely circulate via multiple *Rhinolophus* species. Our current understanding of the geographic distribution of the SARS-CoV and SARS-CoV-2 lineages^11^ possibly reflects a lack of sampling in Southeast Asia, or at least along the Greater Mekong Subregion, which encompasses Myanmar, Laos, Thailand, Cambodia and Vietnam, as well as the Yunnan and Guanxi provinces of China, linking the sampling area of the closest viruses to SARS-CoV-2 identified to date. Finally, pangolins, as well as members of order *Carnivora,* especially the *Viverridae*^5^, *Mustelidae*^6^, and *Felidae*^7^ families are readily susceptible to SARS-CoV-2 infection, and might represent intermediary hosts for transmission to humans. Viruses of the SARS-CoV-2 sublineage, with one exhibiting strong sequence similarity to SARS-CoV-2 in the RBD, were recently detected in distinct groups of pangolins seized during anti-smuggling operations in southeast China^5^. While it is not possible to know where these animals became infected, it is important to note that the natural geographic range of the pangolin species involved (*Manis javanica*) also corresponds to Southeast Asia and not China.

Southeast Asia, which hosts a high diversity of wildlife and where exists extensive trade in and human contact with wild hosts of SARS-like coronaviruses, may represent a key area to consider in the ongoing search for the origins of SARS-CoV-2 ^16^, and certainly in broader coronavirus surveillance efforts. The region is undergoing dramatic land-use changes such as infrastructure development, urban development, and agricultural expansion, that can increase contacts between bats and humans. Continued surveillance of bats and other key wild animals in Southeast Asia is thus crucial, not only to find the SARS-CoV-2 reservoir, but to also be better prepared for the next pandemic.

## Methods

### Ethics statement

The study was approved by the General Directorate of Animal Health and Production and Forest Administration department of the Ministry of Agriculture Forestry and Fisheries in Cambodia. Sampling was conducted under a University of California, Davis Institutional Animal Care and Use Committee approved protocol (UC Davis IACUC Protocol No. 19300). The bat capture and sampling in 2010 was authorized by UNESCO and the National Authority of Preah Vihear.

### Sampling

Testing was performed on archived samples from several programs and field missions (Supplementary Table 1). In 2010, the Muséum national d’Histoire naturelle (MNHN, Paris, France) was mandated by UNESCO and the National Authority of Preah Vihear to conduct a mammal survey in northern Cambodia. During this mission, bats were captured using mist nets and harp traps in two provinces, Preah Vihear and Ratanakiri, to compare bat diversity on the two sides of the Mekong River. A site of bat capture located at the border between Preah Vihear and Stung Treng provinces was later identified as from a cave in the Stung Treng province using GPS data. Bats were morphologically identified at the species level by AH and VTT.

More recent sampling efforts were supported by the USAID-funded PREDICT projects which aimed to strengthen global capacity for detection and discovery of viruses with pandemic potential that can move between animals and people. In missions in 2012 to 2018, samples from bats and carnivorans were collected from free-ranging animals, private animal collection, restaurant, or hunted animals in Battambang, Kampong Cham, Mondulkiri, Preah Vihear, Pursat, Ratanakiri and Stung Treng. Mist nets were used to catch bats. Oral and rectal swabs were collected from live animals which were released after sampling.

The samples from the two sampling missions were stored in viral transport medium solution (VTM; containing tryptose phosphate Broth 2.95%, 145 mM of NaCl, 5% gelatin, 54 mM Amphotericin B, 106 U of penicillin-streptomycin per liter, 80 mg of gentamycin per liter [Sigma-Aldrich, Irvine, UK]) and were held in liquid nitrogen in dewars for transport to the Institut Pasteur du Cambodge where they were stored at −80 °C prior to testing.

The samples were selected and tested for SAS-CoV-2 related virus in the current work as a result of an effort to look at previously-collected samples that had not been initially prioritized for testing or not been tested with RT-PCR assays capable of detecting SARS-CoV-2 related viruses due to resource constraints.

The two bats positive for viruses closely related to SARS-CoV-2 and collected during the MNHN mission were morphologically identified as *Rhinolophus shameli* and their taxonomic status were further confirmed by analyzing the sequences of the cytb gene and the subunit 1 of the cytochrome c oxidase gene (CO1) (Supplementary figure 5).

### RNA Extraction and qRT-PCR

RNA from rectal swabs was extracted using QIAamp^®^ Viral RNA kits. The samples were tested with a pan-coronavirus (pan-CoV) hemi-nested RT-PCR^8^ and by a RT-qPCR known to detect sarbecoviruses^9^, including SARS-CoV-2. A large fraction of these samples has been previously tested with another pan-CoV RT-PCR^17^, which does not detect SARS-CoV-2 like viruses. Both oral swab from the same shameli bats described here tested negative for the presence of betaCoV RNA, despite the high proportion of reads matching the coronavirus (23%) in the rectal swab of RshSTT200. Initial viral isolation attempts were unsuccessful but further isolation is being attempted in several bat cell lines.

### Next generation sequencing

Extracted RNA was treated with Turbo DNase (Ambion) followed by purification using SPRI beads (Agencourt RNA clean XP, Beckman Coulter). We used a ribosomal RNA (rRNA) depletion approach based on RNAse H and targeting human rRNA^10^. The RNA from the selective depletion was used for cDNA synthesis using SuperScript IV (Invitrogen) and random primers, followed by second-strand synthesis. Libraries were prepared using a Nextera XT kit (Illumina) and sequenced on an Illumina NextSeq500 (2×75 cycles).

### Genome assembly

Raw reads were trimmed using Trimmomatic v0.39 ^18^ to remove adaptors and low-quality reads. We assembled reads using the metaspades option of SPAdes/3.14.0 ^19^ and megahit v1.2.9 ^20^ with default parameters. Scaffolds were queried against the NCBI non-redundant protein database ^21^ using DIAMOND v2.0.4 ^22^. Among other putative viruses (hits summarized in Supplementary Table 2), the *Sarbecovirus* genomes identified were verified and corrected by iterative mapping using CLC Assembly Cell v5.1.0 (QIAGEN). Aligned reads were manually inspected using Geneious prime v2020.1.2 (2020) (https://www.geneious.com/), and consensus sequences were generated using a minimum of 3X read-depth coverage to make a base call. The genomes are nearly identical, presenting 3 nucleotides difference between them: g12196a; c20040t; and t24572c). We used Ivar^23^ to estimate the frequency of minor variants (iSNV) from the coronavirus reads. Coverage depth and iSNVs are reported in Supplementary Figure 1.

### Dataset

Complete genome sequence data and metadata of representative SARS-like viruses were retrieved from GenBank, ViPR^24^ and GISAID. Sequences were aligned by MAFTT v.7.467 ^25^, and the alignment checked for accuracy using MEGA v7 ^26^. Accession numbers of all 37 sequences are available in Supplementary Table 4. Separate alignments were generated for the main ORFs. The nucleotide similarities shown in SimPlot^27^ analysis were generated by using a Kimura 2 parameter distance model with a 500-nt sliding window moved along the sequence in 50-nt increments.

### Recombination analysis

We used a combination of 6 methods implemented in RDP5^28^ (RDP, GENECONV, MaxChi, Bootscan, SisScan and 3SEQ) to detect potential recombination events, and conservatively considered recombination signal detected by at least 5 methods. The beginning and end of breakpoints identified with RDP5 were used to split the genome into regions for further phylogenetic analysis.

### Phylogenetic analysis

Maximum-likelihood (ML) phylogenies were inferred using IQ-TREE v2.0.6 ^29^ and branch support was calculated using ultrafast bootstrap approximation with 1,000 replicates ^30^. Prior to the tree reconstruction, the ModelFinder application ^31^, as implemented in IQ-TREE, was used to select the best-fitting nucleotide substitution model.

Bayesian phylogenies were inferred using MrBayes v3.2.7 ^32^, using the GTR substitution model. Ten million steps were run and parameters were sampled every 1000 steps.

### Structure modelling

The three-dimensional structure of the RBD of RshSTT200 was modeled using the SWISS-MODEL program^33^, using SARS-CoV-2 (PDB: 6yla.1) structure as it was the best hit for the RshSTT200 amino acid sequence input.

## Data availability

Sequence data that support the findings of this study have been deposited in the European Nucleotide Archive (Samples ERS5578105 and ERS5578106). The consensus sequences of RshSTT182 and RshSTT200 are also available at the GISAID^34^ with accession numbers: EPI_ISL_852604 and EPI_ISL_852605.

## Acknowledgements

We thank the government of Cambodia for permission to conduct this work. We thank also General Directorate of Animal Health and Production, Department of Wildlife and Biodiversity, Forestry Administration, Ministry of Agriculture, Forestry and Fisheries, Communicable Disease Control Department, Ministry of Health, the Wildlife Conservation Society teams and all students who helped collecting field samples. We extend our gratitude to the Virology Unit team at Institut Pasteur du Cambodge for technical support in laboratory diagnostic. We are grateful to all researchers who have kindly shared genome data on the International Nucleotide Sequence Database Collaboration or on the GISAID.

---

**Supplementary table 1**. Distribution of positive samples within the species for retrospective analyzed samples. Sarbecovirus were only detected in *Rhinolophulus shameli* bat species samples.

**Supplementary table 2**. Viral scaffolds identified in this study

**Supplementary table 3**. Sequence identity for SARS-CoV-2 compared with RshSTT182/RshSTT200 and representative BetaCoV genomes.

**Supplementary table 4**. List of the sequences used for phylogenetic studies.

**Supplementary table 5**. GISAID acknowledgement table

**Supplementary Figure 1.**
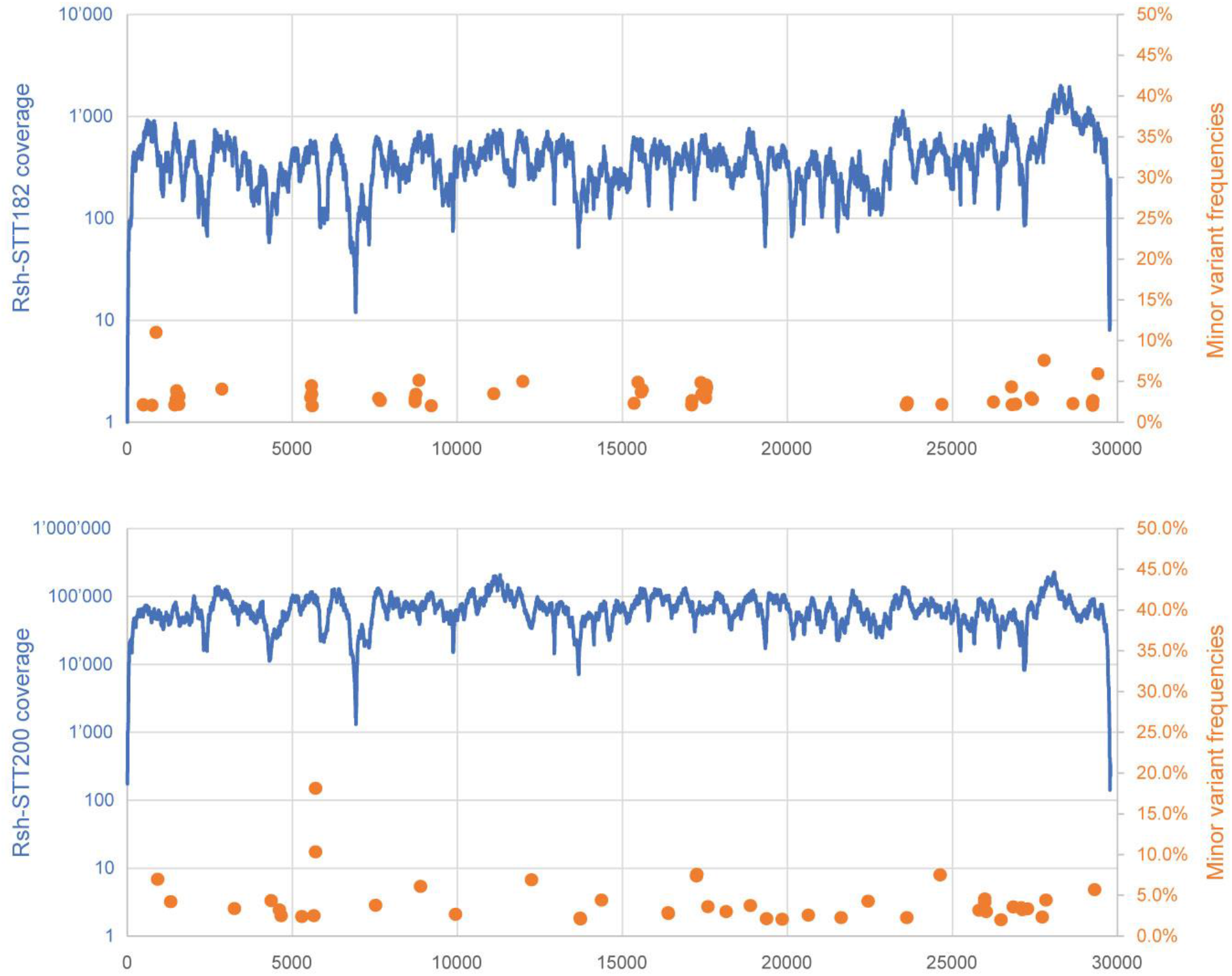
Reads coverage for both IPC viruses and frequency of minor variants for both coronaviruses from Cambodia.

**Supplementary Figure 2.**
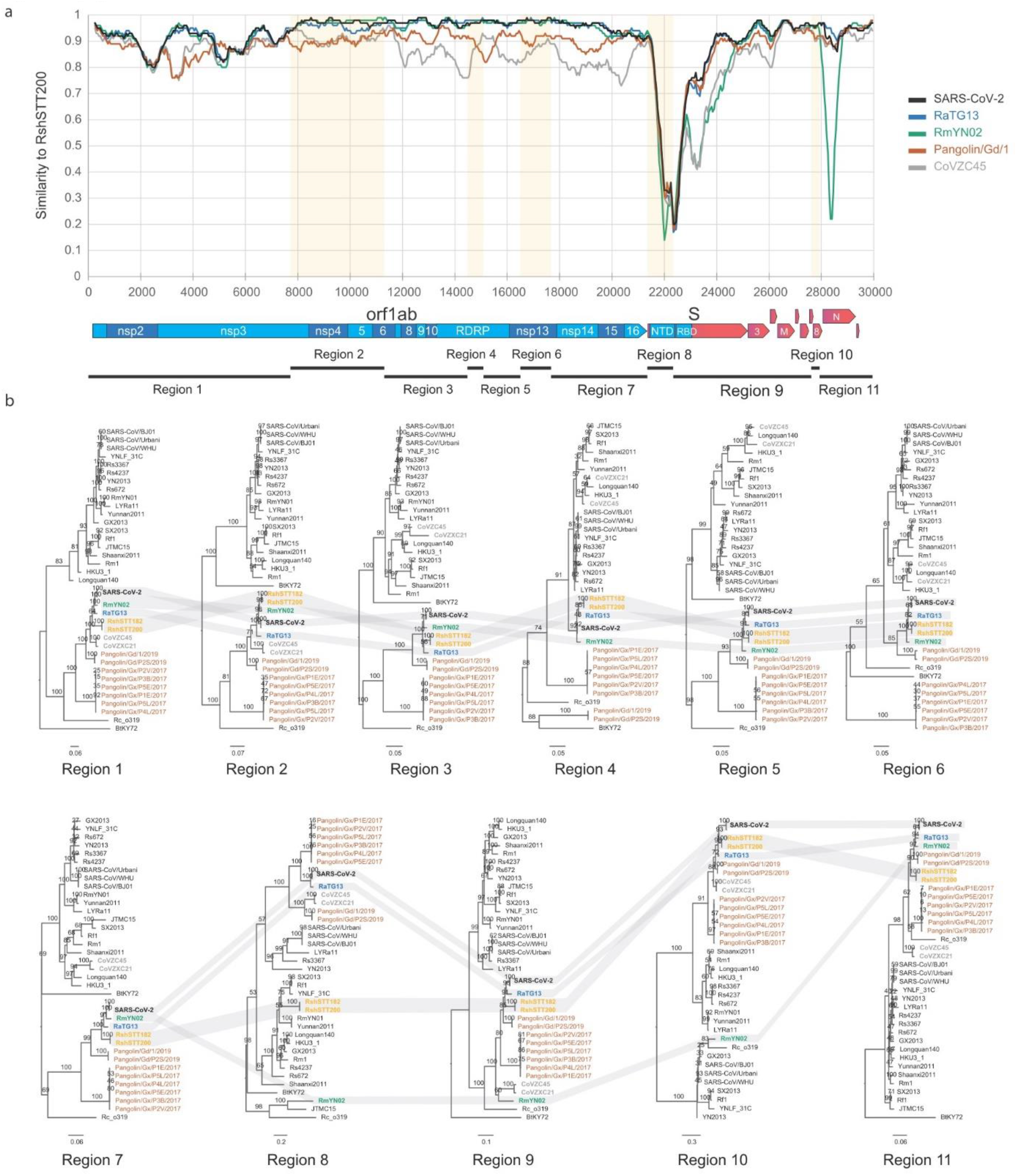
Recombination analysis. a. Sliding window analysis of changing patterns of sequence similarity between RshSTT200 and related coronaviruses from China and Cambodia. ZXC21 and ZC45 were merged for this analysis. The grey shaded boxes indicate regions of RshSTT182/RshSTT200 genomes identified as recombinant. These potential breakpoints subdivide the genomes into 11 regions, indicated by the black bars at the bottom of the similarity plot. The genome organization with the predicted ORFs is shown. b. Phylogenetic tree of genomic regions defined by the recombination analysis. Branch support obtained from 1,000 bootstrap replicates are indicated. SARS-CoV-2 sequences are collapsed, and trees are midpoint rooted for clarity.

**Supplementary Figure 3.**
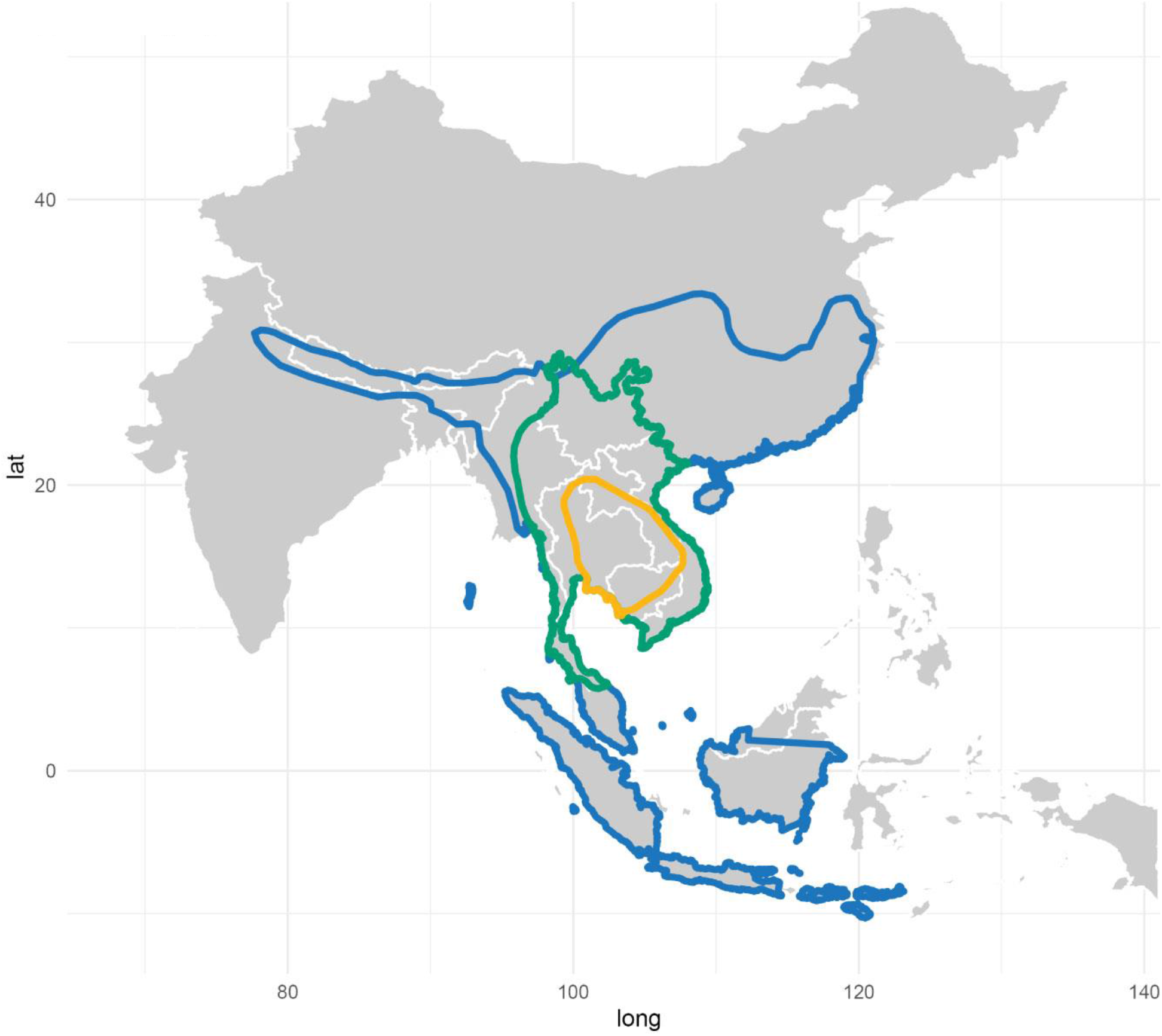
Bats species distribution in South East Asia related to SARS-CoV-2-like virus detection. The distribution of *R. shameli*^35^, *R malayanus*^36^ and *R. affinis*^37^ are shown in orange, green, and blue, respectively. Data retrieved from The IUCN Red List of Threatened Species (https://www.iucnredlist.org/).

